# JARID2 facilitates transcriptional reprogramming in glioblastoma in response to standard treatment

**DOI:** 10.1101/649400

**Authors:** Nora Rippaus, Alexander F-Bruns, Georgette Tanner, Claire Taylor, Alastair Droop, Anke Brüning-Richardson, Matthew A. Care, Joseph Wilkinson, Michael D. Jenkinson, Andrew Brodbelt, Aruna Chakrabarty, Azzam Ismail, Susan Short, Lucy F. Stead

## Abstract

**Background:** Glioblastoma (GBM) is a fatal and incurable brain cancer with a dismal prognosis. In order to impact on this disease, we need to understand how infiltrating, non resectable tumour cells resist chemoradiation and facilitate disease recurrence. To this end, we generated or acquired bulk tumour RNA sequencing data from 45 paired primary and locally recurrent GBM tumours (split into original and validation cohorts) from patients that received standard treatment. We also generated DNA methylation profiles for 9 pairs and sequenced RNA from single cells isolated from a patient derived GBM spheroid model at different timepoints following *in vitro* chemoradiation.

**Results:** We have identified a set of genes with Jumonji and AT-Rich Interacting Domain 2 (JARID2) binding sites in their promoters that are universally dysregulated in post-standard treatment recurrent GBMs compared to the primary tumour. The direction of dysregulation is patient-dependent and not associated with differential promoter DNA methylation. Our *in vitro* experiments suggest that this dysregulation occurs dynamically following treatment as opposed to resulting from selection of cells with specific expression profiles.

**Conclusion:** JARID2 is an accessory protein to a chromatin remodeling complex, responsible for histone modifications observed during cell state transitions in both normal brain and GBM. We propose that JARID2 facilitates GBM recurrence following treatment by indirect transcriptional reprogramming of surviving cells in whichever manner is needed to reproduce the phenotypic heterogeneity required for tumour regrowth *in vivo*. The mechanism of this reprogramming may present a therapeutic vulnerability for more effective treatment of GBM.

## BACKGROUND

Glioblastoma (GBM) is arguably one of the most challenging cancers to treat and is associated with very poor prognosis. This is in part because GBM cells infiltrate the surrounding normal brain making complete surgical removal impossible and, despite subsequent chemoradiation, the remaining cells resist treatment and facilitate tumour regrowth in almost 100% of cases. If we ever hope to more effectively treat GBM we must understand how and why unresected cells resist treatment and form a recurrent tumour. To this end, we and others have focused our attention on molecular profiling of paired primary and recurrent GBM tumours to specifically identify features which are expanded post-treatment and may offer insight into the properties of cells which survive, or the mechanisms that enable their continued proliferation^[1–5]^. As part of the Glioma Longitudinal AnalySiS consortium, we have analysed the genomes of more than 200 paired gliomas and determined that there is no clear evidence for therapy-driven selection of cells bearing specific resistance-conferring mutations (manuscript under review) in agreement with the work of Körber et al.^[3, 6]^. We have, therefore, focused our continued efforts herein on transcriptional features and the possibility of therapy-driven selection of GBM cell populations defined by expression profiles. Transcriptional heterogeneity is evident in GBM: expression profiles align with neurodevelopmental-like hierarchies that span genomic subclones and are functionally distinct, including with respect to treatment sensitivity *in vitro*^[7, 8]^. We generated or acquired RNAseq data from paired primary and recurrent GBMs from a cohort of 23 patients (our original cohort) that underwent standard treatment (debulking surgery followed by chemoradiation with the alkylating drug temozolomide) and had a local recurrence. We then acquired data from an additional 22 such patients (our validation cohort). Our analyses of these data, and of DNA methylation profiles from 9 pairs, consistently highlight the likely role of a Polycomb Repressive Complex 2 (PRC2) accessory protein called JARID2 (Jumonji and AT-Rich Interacting Domain 2) in the transcriptional changes observed after treatment in GBM, via histone modifications and chromatin remodeling. However, the direction of fold change is not consistent across patients. We then performed single cell RNAseq on a patient derived GBM model at different time points following administration of clinically relevant doses of chemoradiation *in vitro* and found that JARID2 associated transcriptional changes occur dynamically after treatment as opposed to resulting from selection of cells with a specific expression profile. JARID2’s interaction with PRC2 is fundamental to specifying neurodevelopmental cell lineages in response to environmental cues, and PRC2 has been shown to be necessary for determining GBM cell phenotypes based on tumour microenvironmental pressures, though the role of JARID2 in this has never been investigated^[9–12]^. We propose that GBM recurrence results from JARID2-associated transcriptional reprogramming, via PRC2, of unresected cells in whichever direction enables recapitulation of the transcriptional heterogeneity needed for continued tumour growth *in vivo*^[13]^. Therapeutic targeting of the mechanism of such reprogramming may constitute a more effective treatment strategy than targeting of the cell types that lie either side of the interconversions.

## RESULTS

### Differential expression indicates a therapy-driven shift in neurodevelopmental genes

Genes that were differentially expressed (DE) in the recurrent versus primary GBMs in our original cohort (Supp.Table.1) were significantly enriched for those involved in cell development and lineage determination, specifically in relation to neurodevelopment (Fig.1a and Supp.Table.2). Brain cell fate is orchestrated by the combined actions of transcription factors (TFs) and chromatin remodeling complexes, both of which have been implicated in establishing functionally heterogeneous transcriptional hierarchies in gliomas^[14, 15]^. We therefore reasoned that specific DNA-binding factors may coordinately regulate the genes we observe to be altered after treatment, potentially highlighting certain cell types that survive. Candidate master transcriptional regulators can be identified from expression data via gene set enrichment analysis (GSEA). However, we found that many neurodevelopmental TFs were missing from publicly available gene sets, so we first developed a more comprehensive DNA-binding factor gene set using ChIPseq data from the Gene Transcription Regulation Database (see Methods)^[16]^.

**Fig. 1.**
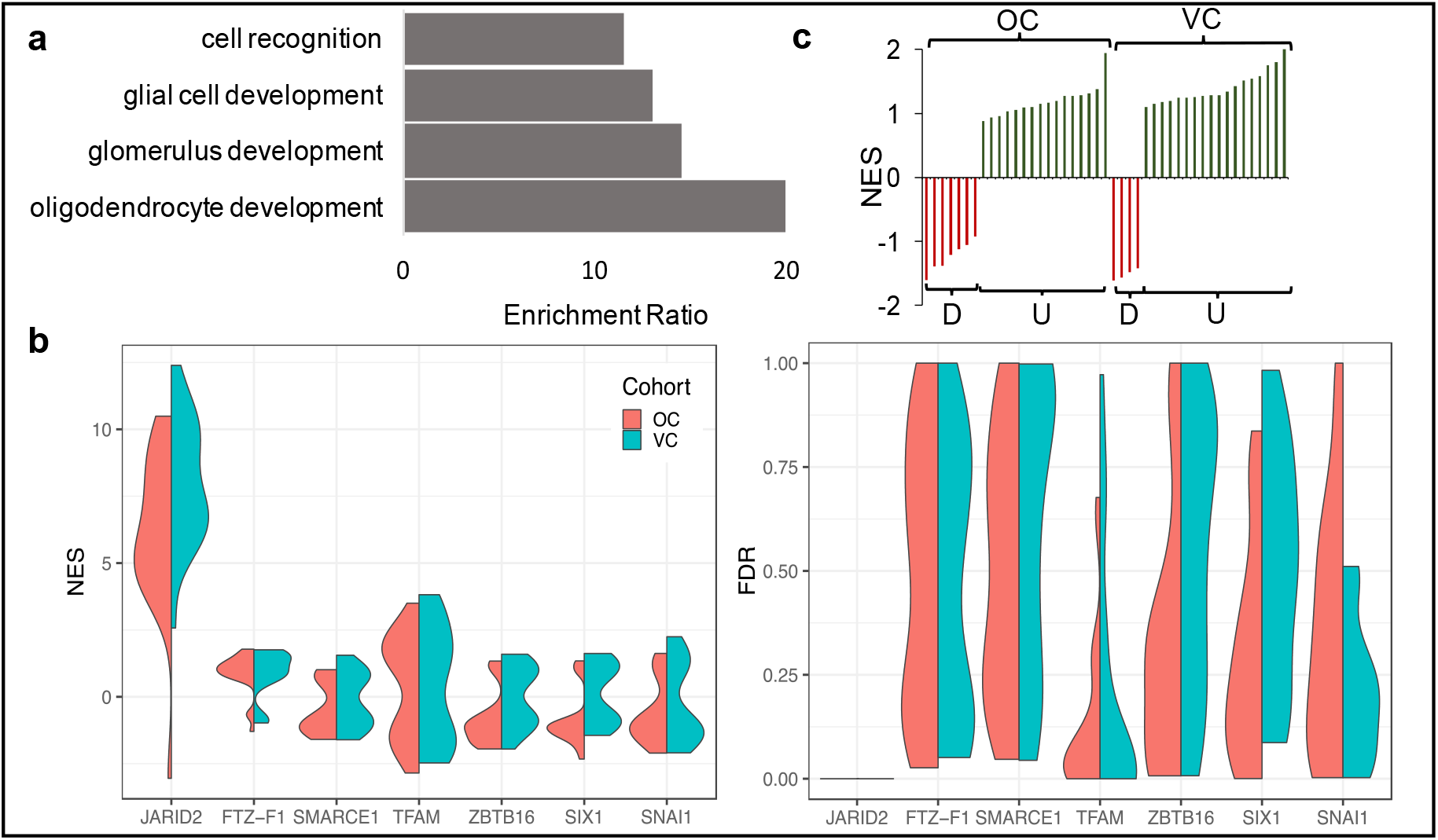
**a)** Biological processes enriched in the genes differentially expressed between matched primary and recurrent GBMs (enrichment ratio and term count>10, p<0.0005); **b)** Per-patient normalised enrichment scores (NES, left plot) and false discovery rates (FDR, right plot) for the top-scoring promoter-binding factors associated with gene expression changes in recurrent vs primary GBMs, highlighting the significance of JARID2; **c)** The NES for JARID2 for each patient (x-axis) when direction of fold change is taken into account, shows that there are two response subtypes based on whether genes are up (U) or down (D) regulated. OC: original cohort. VC: validation cohort.

### Genes with JARID2 binding sites in their promoters (JBSgenes) are consistently and significantly dysregulated in recurrent versus primary GBMs and stratify patients into two response subtypes

We performed per-patient GSEA, using our novel gene set, with genes pre-ranked by the magnitude of fold change in expression between the primary and recurrent tumour. Genes with a Jumonji and AT-Rich Interacting Domain 2 (JARID2) binding site in their promoters (JBSgenes) were the most significantly, consistently and highly enriched within the genes changed after therapy across patients: the normalized enrichment score (NES) for JARID2 was significant in all patients (FDR<0.05) and gave the highest score in 91% (n=21/23) (Fig.1b). To determine whether JBSgenes were consistently up-regulated or down-regulated after treatment, we repeated the analysis including the direction of fold change in the gene ranking. We found that the JBSgenes were altered in a consistent direction per patient but across patients the direction varied: in 30% (n=7/23) the JARID2 enrichment was being driven by down-regulation of JBSgenes (hereon referred to as D response subtype) and in the remaining 70% (n=16/23) it through up-regulation (U response subtype) (Fig.1c).

We acquired data from an additional 22 paired primary and recurrent GBMs from patients who underwent standard treatment and who had a local recurrence (the validation cohort) which corroborated our findings (Figs.1b and c) with a similar ratio of D response subtype (n=4/22) and U response subtype (n=18/22) patients (Fisher’s exact test, p=0.52)^[5]^.

### The same JBSgenes are dysregulated in each response subtype and their promoter DNA is unmethylated in both primary and recurrent GBMs

To investigate whether the JBSgenes driving the enrichment differed across individual patients or between response subtypes, we quantified how often each gene was present in the leading edge of the GSEA results across the 45 patients in the original and validation cohorts combined.

335 genes were observed in the leading edge of more than 50% of patients (denoted LE50 genes) and 43 genes in more than 70% (LE70 genes) (Supp.Table.3). The per-patient fold-changes of the LE50 genes showed that these same JBSgenes drive the enrichment across patients irrespective of response subtype i.e. the same genes are downregulated in D response patients as are upregulated in U response patients (Fig.2a). The LE50 genes are enriched in functional annotations associated with neurodevelopment and neuronal differentiation such as synaptic plasticity and interneuronal communication (Fig.2b and Supp.Table.4). To investigate the DNA methylation status of the promoters of these genes in primary and recurrent GBMs, in comparison to other genes, we performed genome-wide methylation arrays on DNA from 9 pairs (see Supp.Table.1). No gene promoters were differentially methylated between primary and recurrent samples (q>0.45 for all genes). However, the distribution of promoter methylation across all genes was significantly different to that of the JBSgenes and the LE50 genes in isolation, revealing that the DNA in the promoters of the latter two is unmethylated in both the primary and recurrent GBMs (Fig.2c). This indicates that the change in expression of these genes that we observe post-treatment is not driven by DNA methylation.

**Fig.2.**
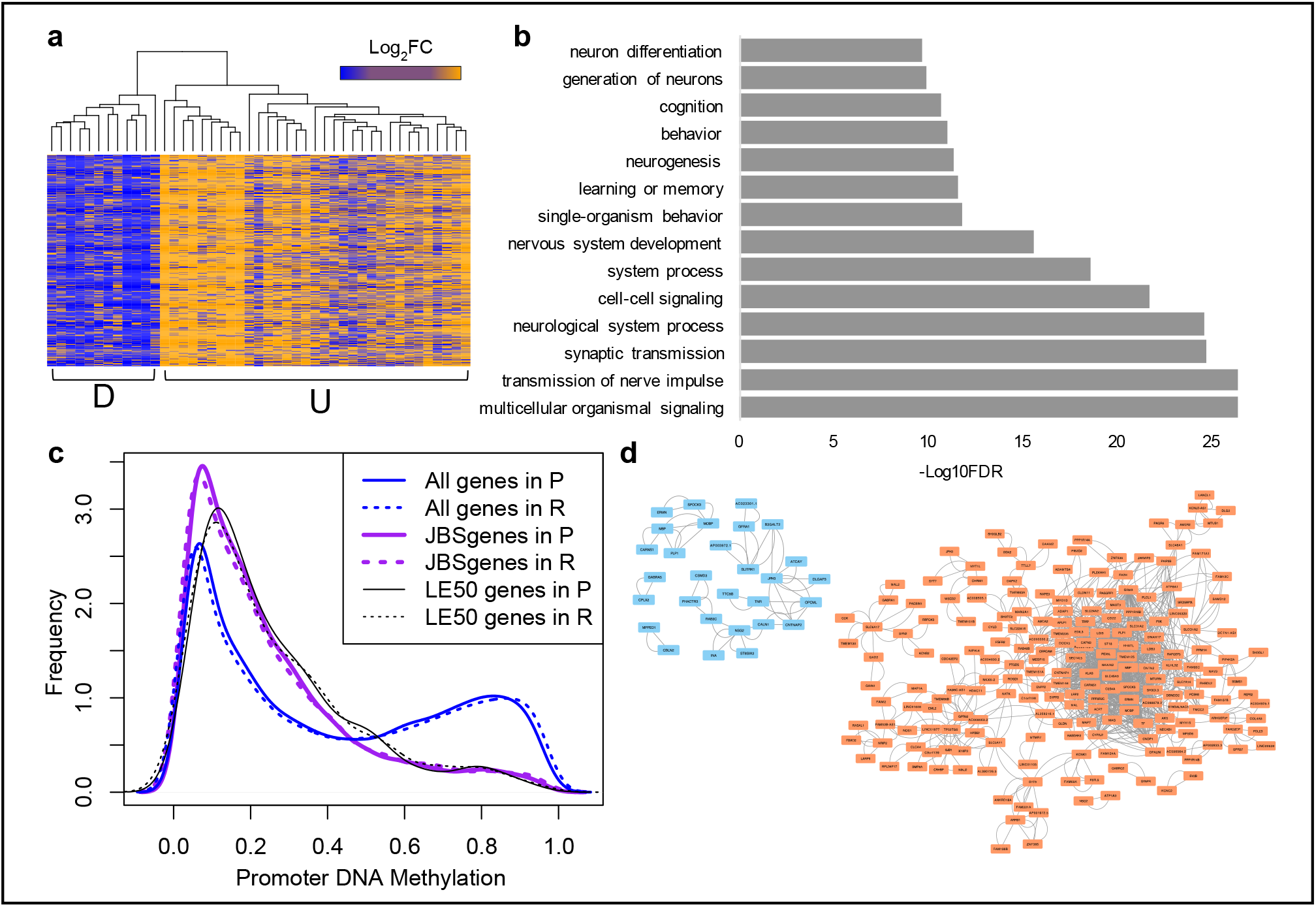
**a)** Heatmap of fold change in expression after treatment per patient (columns) for the genes in the leading edge of the JARID2 GSEA results in more than 50% of patients across both cohorts (LE50 genes, rows). These same genes are upregulated in U response subtype patients as are downregulated in D response subtype patients; **b)** The biological processes (with <2000 terms) most enriched in LE50 genes; **c)** The distribution of average promoter DNA methylation for all genes, JARID2 binding site (JBS)genes and LE50 genes in primary (P) and recurrent (R) GBM samples; **d)** Networks of genes (nodes) for which expression is highly correlated (edges: R>|0.9|) with LE70 genes in the primary (blue, top left) and recurrent (orange, right) GBM samples.

### LE70 JBSgenes are more coordinately expressed in recurrent GBM

We hypothesised that JARID2 is involved in the tighter co-regulation of leading edge JBSgenes in GBM tumours after treatment, independent of whether that results in an increase or decrease in their expression. To investigate this, we identified all genes for which expression is highly correlated (R>|0.9|) in the primary GBMs and recurrent GBMs separately. We then determined the prevalence of the LE70 genes and their connectivity in these two correlation networks. We found that the LE70 JBSgenes correlate with significantly more genes and with significantly more connectivity in recurrent versus primary samples: 1% (29/2603) in the primary GBM network compared to 7% (202/2855) in the recurrent GBM network (chi-squared; p=0 for both tests) (Fig.2d). This implies that either cells in which these genes are co-regulated by JARID2 become more prevalent post-treatment, or that their co-regulation by JARID2 occurs in response to treatment. To inspect this further we designed an *in vitro* experiment to investigate the time course of JBSgene dysregulation following treatment.

### Single cell analysis indicates that JARID2 associated gene co-regulation is an adaptive response to therapy

We cultured two plates of spheroids directly from a freshly resected primary GBM, in serum-free conditions. We treated one plate with physiologically relevant single doses of TMZ (30μM) and radiation (2Gy). We captured and sequenced RNA from single cells from spheroids one week post-treatment when there was a significant deviation in the untreated vs treated spheroid growth curves, and three weeks post-treatment when growth of the treated spheroids appeared to have recovered (Fig.3). Our bespoke GSEA revealed that JBSgenes were significantly enriched amongst the genes altered in treated versus untreated spheroids three-weeks post-treatment (FDR=0.18) but not one week post-treatment (FDR=0.65). Furthermore, the genes that were DE (p<0.05) between treated and untreated cells included significantly more LE50 genes at the three-week time point compared to the one week time-point (chi-squared, p=0.007). These results suggest that the universal JBSgene dysregulation that we observe in recurrent tumours is not caused by selection of a fixed transcriptional profile, but rather transcriptional reprogramming following treatment.

**Fig. 3.**
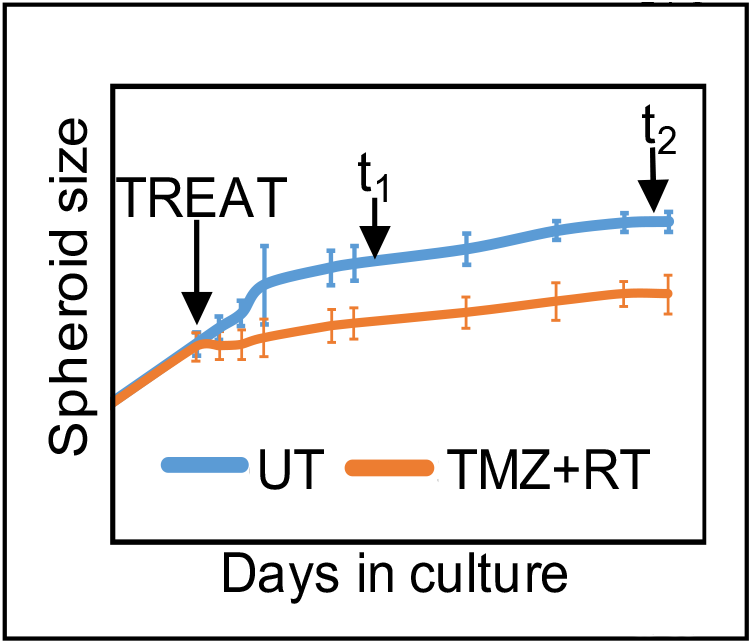
Growth curves for untreated (UT) and treated (TMZ+RT) patient-derived GBM spheroids. Time of treatment (TREAT) is indicated (arrow), in relation to single cell capture 1 week (t_1_) and 3 weeks (t_2_) post-treatment.

### JARID2 is involved in cell plasticity and implicated in interconversions between cell states in glioma

JARID2 is an accessory protein responsible for the genomic positioning of Polycomb Repressive Complex 2 (PRC2)^[12]^. PRC2 is a chromatin remodeller that is indispensable for lineage determination during neurogenesis^[10]^. It is responsible for the trimethylation of lysine 27 on histone H3 (H3K27me3) which results in epigenetic silencing of the marked gene. PRC2 is directly implicated in cell plasticity in GBM by studies showing that its catalytic subunit is required to enable conversions between stem-like and differentiated cell types^[11]^. It is also indirectly implicated by the fact that the prevalence and location of H3K27me3 significantly differs between normal brain and glioma cells, and between GBM cells with different phenotypes^[17]^. These differences occur most frequently at bivalent promoters: those harbouring both the repressive H3K27me3 and an activating mark (H3K4me3) causing the gene to be silenced but primed for activation upon PRC2 disassociation and H3K27 demethylation. Developmental gene promoters are commonly bivalent in embryonic stem cells to enable subsequent rapid activation of specific lineage determination genes once cell fate is resolved, further highlighting PRC2’s role in cell-type transitions^[18]^.

To determine whether JBSgenes, and specifically the LE50 genes, are implicated in cell type switching we mined published data on histone marks (H3K27me3 and H3K4me3) and gene expression in different normal and GBM cell types ^[17, 19]^. Normal brain cell types were human neural stem cells (NSCs) and normal human astrocytes (NHAs). Different GBM ‘cell types’ pertains to the ability to derive phenotypically distinct cell lines from the same patient GBM under different conditions: those which enrich for glioma stem cells (GSCs, which can be considered analogous to NSCs in the normal brain) and those which enrich for differentiated glioma cells (DGCs, somewhat analogous to NHAs)^[20]^. We first quantified promoter status for each different cell types: active=H3K4me3; repressed=H3K27me3; bivalent=H3K27me3+H3K4me3; and neither mark. In support of the role of JARID2 as a PRC2 accessory protein, and increasing confidence that our novel gene set has captured bona fide JBSgenes, we found that H3K27me3 was significantly enriched at JBSgenes in all cell types investigated (Fig.4: middle compared to top barplot in each panel; grey and orange shading pertains to promoters with the H3K27me3 mark), and that this was particularly pronounced at bivalent promoters (Fig.4; orange shading). Moreover, the presence of H3K27me3, again especially at bivalent promoters, was further significantly enriched within the LE50 subset of JBSgenes (Fig.4 bottom compared to middle barplot in each panel). We then characterized changes in promoter status between cells of different phenotype in the normal brain (NPC vs NHA) and GBM (GSC vs DGC: Fig.5). We found, in agreement with the results from the original publication, that changes involving H3K27me3 are more pronounced than any other. We further found that this is significantly more evident in the promoters of JBSgenes: 85% of changes between NPC and NHA involve H3K27me3 at JBSgene promoters compared to 76% at all gene promoters (chi-squared, p= 3.9×10^−8^) and 93% of changes between GSC and DGC involve H3K27me3 at JBSgene promoters compared to 78% at all gene promoters (Fig.5; chi-squared, p= 5.6×10^−4^). This implicates JARID2 in chromatin remodeling of gene promoters that differ between cell types in the both normal brain and in GBM.

**Fig. 4.**
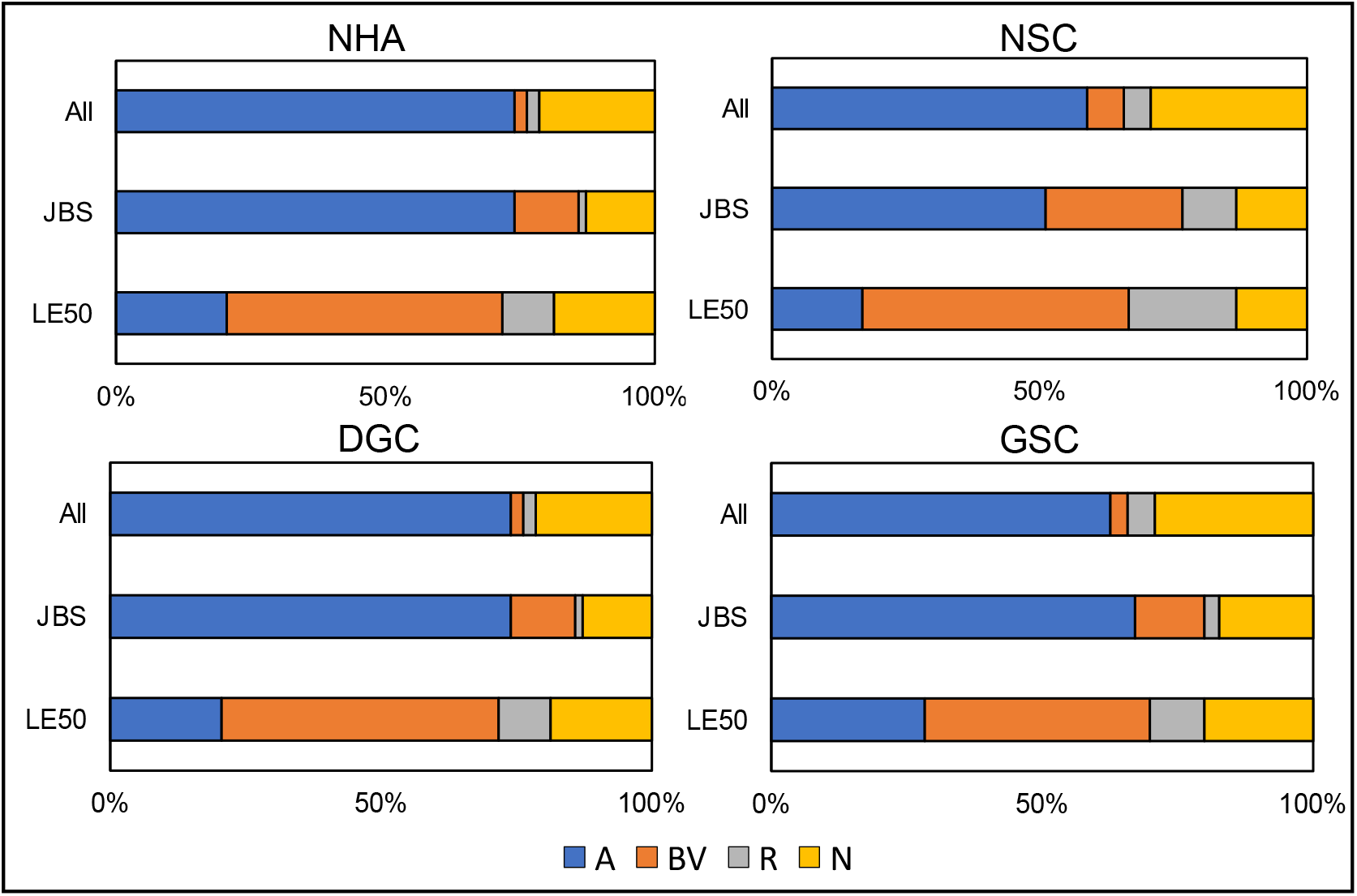
Relative quantification of promoter status, with respect to specific histone marks, across all genes compared to JBSgenes and LE50 genes in normal human astrocytes (NHA), neural stem cells (NSC), differentiated glioma cells (DGC) and glioma stem cells (GSC). A=active=H3K4me3; R=repressed=H3K27me3; BV=bivalent=H3K27me3+H3K4me3; and N=neither mark.

**Fig.5.**
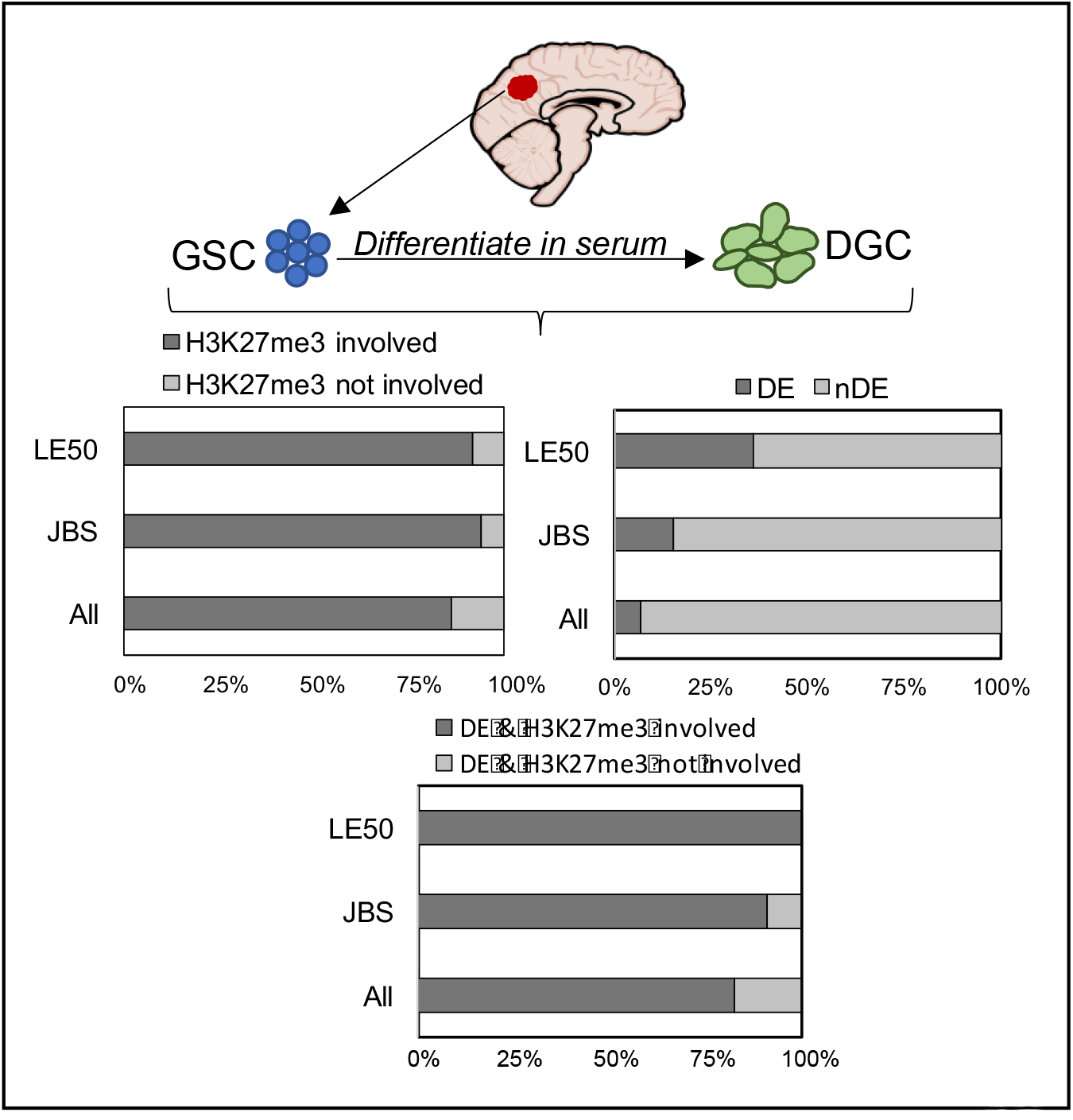
Glioma stem cell (GSC) cell lines can be derived from patient GBM tumours and then further cultured in serum to form differentiated glioma cells (DGC). The bar charts indicate the proportion of genes for which H3K27me3 is involved in any changes in promoter status (top left), or which were differentially expressed (DE, top right), or both (bottom), between matched GSC and DGC cell lines for all genes, JBSgenes or LE50 genes separately.

To determine whether this remodeling associates with gene expression changes we identified genes that are DE (q<0.2) between GSC and DGC using RNAseq data from Patel et al.^[19]^ from matched lines derived from three patient samples. As shown in Fig.5, we found that significantly more JBSgenes are DE (15% JBSgenes vs 5% of non-JBSgenes; chi-squared, p=0) and that, within the JBSgenes, significantly more LE50 genes are DE (36% of LE50 genes vs 15% of remaining JBSgenes; chi-squared, p=2.5×10^−5^). We then overlaid the expression and histone mark data and found, as also shown in Fig.5, that chromatin remodeling involving H3K27me3 is more pronounced at DE JBSgenes (91% of changes at DE JBSgene promoters involve H3K27me3) and DE LE50 genes (100% of changes) compared with all DE genes (where 82% of promoter status changes involve H3K27me3). Together, these data suggest that chromatin remodeling associated with JARID2 affects the expression of genes that distinguish different cell types in the normal and human brain.

## DISCUSSION

We found that the genes differentially expressed between pairs of primary and locally recurrent GBM tumours post standard treatment were enriched for those involved in brain cell development and lineage determination, suggesting that GBM cell types defined within neurodevelopmental-like transcriptional hierarchies may be associated with treatment resistance and tumour regrowth in patients. Brain and GBM cell specification results from a combination of the concerted action of transcription factors (TFs) and differential genome accessibility imposed by a variety of chromatin remodeling molecules and complexes. We, therefore, applied an unbiased approach to investigate whether any such DNA-binding factors were repeatedly implicated across patients in the master regulation of genes for which we observe altered expression in recurrent versus primary GBM tumours. We found that genes with Jumonji and AT-Rich Interacting Domain 2 (JARID2) binding site(s) in their promoter (JBSgenes) are consistently and significantly dysregulated in recurrent GBM tumours, in both our original and validation cohorts. JARID2 is indirectly responsible for eliciting the programmes of epigenetic gene silencing required for cell lineage determination during neurodevelopment^[9]^. It does so by docking Polycomb Repressive Complex 2 (PRC2) to specific genomic loci where it trimethylates H3K27 to repress gene expression^[12, 21]^. As well as normal brain cell delineation, PRC2 and the H3K27me3 mark have been specifically implicated in cell state transitions in glioblastoma^[11, 17, 22]^. However, the involvement of JARID2 in this process, and the genes in which expression is altered by this mechanism to dictate GBM cell phenotype have not previously been elucidated. Our results highlight subsets of genes with JARID2 binding sites in their promoters (JBSgenes) that are most commonly dysregulated (LE50 in more than 50% and LE70 in more than 70% of patients) in GBM tumours after treatment. In support of the notion that expression of these genes is regulated by JARID2-assoicated mechanisms in GBM, we have shown that the DNA in their promoters is unmethylated in both primary and recurrent samples and that their promoters are significantly more associated with the H3K27me3 mark than those of other genes in GBM cell lines. The increasing importance of this coordinated regulation by JARID2 in GBM after treatment is indicated by the larger and more connected networks of highly correlated LE70 JBSgenes in recurrent versus primary samples.

Together these results could suggest that specific GBM cell types, defined by transcriptional profiles resulting from JARID2-associated epigenetic programming, resist treatment and expand during tumour recurrence. It is widely thought, for example, that glioma stem cells specifically resist treatment and are responsible for GBM regrowth^[23, 24]^. However, a confounding result in relation to this interpretation is the fact that, whilst a specific subset of JBSgenes are universally dysregulated in recurrent versus primary tumours, the direction of dysregulation is inconsistent; the same genes are upregulated in the recurrence in ~70% of patients and down-regulated in the remaining 30%. Furthermore, our *in vitro* work indicates that changes in expression of JBSgenes occurs dynamically following treatment as opposed to resulting from an increased signal from expansion of a fixed transcriptional profile. Our hypothesis is, therefore, that JARID2-associated chromatin remodeling is not a treatment resistance mechanism per se, but a mechanism by which GBM tumours recover from treatment to enable regrowth. In this way, the different directions of gene dysregulation are owing to the need to recapitulate the GBM transcriptional heterogeneity, required for tumour growth *in vivo*, from whichever cell types survived in that particular patient^[13, 25, 26]^. This is supported by our findings that the JBSgenes dysregulated during treatment are a) significantly more likely to have bivalent promoters (i.e. poised for activity relating to lineage decisions in response to environmental queues) and b) significantly enriched amongst the genes differentially expressed between cells at either end of the GBM transcriptional hierarchy (i.e. glioma stems cells and differentiated glioma cells). Recent landmark findings also support our hypothesis, which posits that different GBM cell types are able to resist treatment and that interconversions between cell types, as opposed to one-way transitions down a differentiation pathway, are needed to enable tumour regrowth: i) differentiated GBM cells form networks *in vivo* that enable them to better survive chemoradiation, negating the idea that only stem-like cells are able to survive^[27]^; ii) glioma stem-cells, needed for tumour regrowth owing to their proliferation capabilities, can form via non-hierarchical conversion of differentiated cells in GBM^[11, 14, 25]^.

Our hypothesis represents a paradigm shift, also recently suggested by Dirkse et al.^[25]^, that challenges the notion that effective treatment of GBM will be possible by therapeutically targeting any one cell population, such as glioma stem cells. Instead, we propose that effective treatment will only be possible by targeting the mechanisms of GBM cell plasticity resulting from transcriptional reprogramming, which our results suggest are fundamentally linked with the role of JARID2. The precise nature of this role, and its potential for therapeutic targeting, are the focus of our ongoing work.

## CONCLUSION

We have found that a subset of genes is universally dysregulated in patient GBMs following standard treatment, likely because of epigenetic remodeling of their promoters via mechanisms involving JARID2, as an adaptive response that facilitates tumour regrowth. The direction of this adaptive response is, however, not constant across patients and may depend upon the cell state transitions needed to recapitulate transcriptional heterogeneity in the recurrent tumour. This is the first time that JARID2 has been implicated in GBM cell plasticity in association with tumour recurrence, and highlights subsets of genes that may be involved in cell state transitions required for adaption of GBM tumours to treatment.

## METHODS

### Archival Samples and Profiling Data

Four independent sources of paired patient GBM samples (surgical tissue from primary GBM and subsequent recurrent samples) were used in this work. Samples were allocated to the original cohort if they had undergone whole transcriptome RNA sequencing, and to the validation cohort if they had undergone poly-A transcriptome sequencing. Clinical information and cohort assignment are given in Supp.Table.1.

#### Stead Samples

21 patients from four tissue banks (Leeds, Liverpool, Cambridge and Preston) with tumour in paraffin blocks. Ethical approval was acquired (REC 13/SC/0509). RNA and DNA was extracted from the same cells from neuropathologist annotated tumour regions (>60% cancer cells) using appropriate Qiagen kits (Qiagen, Sussex, UK). Paired end strand-specific whole transcriptome libraries were prepared for 16 pairs using the NEBNext Ultra Directional RNA Library Prep Kit for Illumina (New England BioLabs, UK), following rRNA depletion with NEBNext rRNA Depletion Kit or Ribo-Zero Gold. Libraries were sequenced on an Illumina HiSeq. DNA from 9 pairs (4 of which also underwent RNAseq) was profiled using the Illumina Infinium Human Methylation 450K Bead Chip array.

#### Rabadan Samples

Nine patients from Wang et al.^[4]^ with transcriptome sequencing data (7 with whole transcriptome data and 2 with poly-A transcriptome data) for paired tumours, downloadable from the sequencing read archive (accession SRP074425).

#### Verhaak Samples

Four patients from Kim et al.^[1]^ with poly-A transcriptome sequencing alignment data acquired, and converted to raw fastq format, following application to the European Genome-Phenome Archive (accession EGAS00001001033).

#### Nam Samples

16 patients from Kim et al.^[5]^ with poly-A transcriptome sequencing alignment data acquired, and converted to raw fastq format, following application to the European Genome-Phenome Archive (accession EGAD00001001424).

### Sequencing Data: Alignment, Differential Expression, Functional Enrichment and Correlation Analysis

RNAseq data was analysed as previously described except that reads were aligned to human reference genome GRCh38, using the gencode.v27 genome annotation as a guide, using STARv2.4.3a and functional enrichment analysis was done using WebGestalt ^[28–30]^.

### Gene set enrichment analysis (GSEA)

We developed a novel gene set file for use in GSEA using the Gene Transcription Regulation Database (GTRD v18.01), which contained the genomic binding locations of 682 human DNA-binding factors from 4236 chromatin immunoprecipitation sequencing (ChIPseq) experiments^[16]^. A gene was assigned to a DNA-binding factor’s gene set if its promoter (transcription start site from gencodev27 ±1kbp) contained a binding site for that factor in ≥2 independent ChIPseq experiments. We first performed pre-ranked GSEA, per patient, ordering genes by the magnitude of fold change in expression log_2_(|recurrent FPKM +0.01/primary FPKM+0.01|) in classical mode. To indicate directionality of dysregulation we then ranked genes by absolute fold changes ie. using log_2_(recurrent FPKM +0.01/primary FPKM+0.01) and weighted by magnitude^[31]^.

### DNA methylation analysis

The RnBeads package was used to import, quality check and preprocess IDAT files and then perform pairwise differential methylation analysis. The combined and adjusted p-value (comb.p.adj.fdr) in the promoter results file was used to determine significance. The average methylation signal for each promoter (mean.mean) was extracted for both the primary and recurrent samples and used to plot the distribution for different genes subsets in R using the density function.

### Patient-derived spheroids

A patient presenting with a suspected GBM was consented for the use of their tissue in research through the Leeds Multidisciplinary Research Tissue Bank (REC 15/YH/0080). GBM diagnosis was confirmed intraoperatively by a neuropathologist who identified a tumour cell rich piece of tissue, surplus to diagnosis, for transport to the laboratory in cold PBS for use in this work. Tissue was washed in PBS and chopped in Accutase (Sigma-Aldrich, 500μL) before incubation at 37°C for 5 min. The sample was triturated and Neural Basal (NB) medium, consisting of Neurobasal Medium, N2 and B27 supplements (ThermoFisher, 250mL, 1.25 and 2.5mL respectively), recombinant basic fibroblast growth factor (bFGF), and epidermal growth factor (EGF) (R&D Systems 40ng/mL each), was added to a total volume of 10 mL prior to spinning (1200 rpm at 5 min). The pellet was resuspended in 5ml DNaseI then 1ml RBC lysis buffer (VWR International) with 1 min incubation at room temp, addition of PBS to 10mL and further spinning following each resuspension. The pellet was resuspended in 10mL PBS, filtered via a 70μm and 30μm strainer consecutively and counted. Finally, cells were resuspended in NB-medium to a concentration of 2×10^4^cells/1mL with 200μL of this cell suspension added into each well of an ultra-low-adherence plate and incubated at 37°C 5%CO2. 100μL of medium was replaced per well every 3 days. Cells were imaged regularly on the EVOS Cell Imaging System (ThermoFisher) until they reached approximately 300μm in diameter. At this point TMZ (Sigma) was concentrated in 100μL of NB media and used in a media replacement for one plate of cells to give a final dose, per well, of 30μM; one hour later the same plate was irradiated with 2Gy.

### Single cell capture and sequencing

Single cells were captured from treated spheroids 1 week and 3 weeks post-treatment, and cells from untreated spheroids the following day, by extracting and combining 8 spheroids per time point and dissociating them via a PBS wash, Accutase incubation, and further PBS washing (as above) to a concentration of 2.5 × 10^5^ cells/mL. Cells were diluted in C1 cell suspension reagent at a ratio of 3:2, respectively. Single cells were captured on a medium (10–17 μM) C1 Single-Cell Auto Prep IFC for mRNAseq, lysed and underwent on-chip cDNA amplification via SMART Seq2 according to the manufacturer’s instructions using the Fluidigm C1 Single-Cell Auto Prep System and protocols on Script Hub™ (Fluidigm)^[32]^. cDNA was quantified using the High Sensitivity DNA Assay on the Agilent 2100 Bioanalyser and paired end Nextera XT (Illumina) libraries were made and sequenced, with multiplexing using indexes provided by Dr Iain Macaulay, on an Illumina HiSeq. RNA sequencing data was processed and expression was quantified as per the bulk tissue. GSEA analysis was performed twice using expression (in fragments per kilobase per million mapped) datasets to identify differences between untreated and treated cells at the 1 week timepoint and three week timepoint separately. Differentially expressed genes were identified using Seurat^[33]^.

### Cell type expression and histone modification status

Data on gene promoter histone status in different normal and GBM cell types were extracted from the supplementary material (Table S2) from Rheinbay et al.^[17]^. This included data from three patient GBM tumours that had been used to derive glioma stem cells (GSCs) which were subsequently cultured in serum to produce differentiated glioma cells (DGC). We required that all three samples had been assigned the same status to be included in our analyses. Raw sequencing data was downloaded on a further three GSC and DGC pairs from Patel et al.^[19]^ via the Gene Expression Omnibus (accession GSE57872). These data were processed and aligned, and pairwise differential expression analysis was performed, exactly as for the paired primary and recurrent RNAseq data. Quantification and assessment of the significance of overlap in genes with different promoter states and/or differential expression with the JBSgenes or LE50 genes was done in the R statistical package.

## LIST OF ABBREVIATIONS

ChIP: Chromatin immunoprecipitation
DE: Differentially expressed
DGC: Differentiated glioma cell
GBM: Glioblastoma
GSC: Glioma stem cell
GSEA: Gene set enrichment analysis
H3K4me3: Trimethylated histone 3 at lysine 4
H3K27me3: Trimethylated histone 3 at lysine 27
JARID2: Jumonji And AT-Rich Interaction Domain Containing 2
JBSgenes: JARID2 binding site genes
LE: Leading edge
NES: Normalized enrichment score
PRC2: Polycomb Repressive Complex 2
RNAseq: RNA sequencing
TF: Transcription factor

## DECLARATIONS

### Ethics approval and consent to participate

All patients whose data was used in this study consented to the use of their tissue in research. The data generated for this study was used in accordance with ethical approval acquired from NHS NRES Committee South Central - Oxford A (REC 13/SC/0509).

### Consent for publication

Not applicable

### Availability of data and materials

The RNAseq datasets analysed during the current study are available in the following repositories: Rabadan samples are downloadable from the sequencing read archive (accession SRP074425); Verhaak and Nam samples are available following application to the European Genome-Phenome Archive (accession numbers EGAS00001001033 and EGAD00001001424, respectively). The RNAseq and DNA methylation datasets generated during the current study are not publicly available as data transfer agreements are required to ensure compliance with ethical approvals but are available from the corresponding author on reasonable request.

### Authors’ contributions

LFS devised the project. LFS and SS acquired funding. MDJ and AB sourced samples and provided clinical annotation. NR and AFB processed samples following annotation and diagnostic confirmation from AC and AI. LFS, GT, AD, MC and JW performed data analysis and LFS interpreted it. NR performed experimental work with assistance from ABR. NR and CT captured and sequenced single cells. LFS wrote the manuscript, which was reviewed and approved by all authors.

## Acknowledgements

Tissue was sourced from: the Brain Tumour Northwest tissue bank (including the Walton research tissue bank) funded by the Sidney Driscol Neuroscience Foundation and part of the Walton Centre and Lancashire Teaching Hospitals NHS Foundation Trusts; the Leeds Multidisciplinary Research Tissue Bank with staff funded by the PPR Foundation and part of the University of Leeds and Leeds Teaching Hospitals NHS Trust; the Human Research Tissue Bank, part of the Cambridge University Hospitals NHS Foundation Trust. Access to the latter was coordinated as part of the UK Brain Archive Information Network (BRAIN UK) which is funded by the Medical Research Council and brainstrust. We would like to thank Dr Iain Macaulay of The Earlham Institute in Norwich for his support for single cell work, and for providing indexes to enable single cell multiplexing. We would also like to thank the authors of publications from which we were able to access data for use in our research.

